# Characterisation of the symbionts in the Mediterranean fruitfly gut

**DOI:** 10.1101/2021.06.09.447743

**Authors:** M. Darrington, P.T. Leftwich, N.A. Holmes, L.A. Friend, N.V.E. Clarke, S.F. Worsley, J.T. Margaritopolous, S.A. Hogenhout, M.I. Hutchings, T Chapman

**Affiliations:** School of Biological Sciences, University of East Anglia, Norwich Research Park, Norwich, NR4 7TJ, UK; Department of Plant Protection, Institute of Industrial and Fodder Crops, Hellenic Agricultural Organization–DEMETER, Volos, Greece; Department of Crop Genetics, John Innes Centre, Norwich Research Park, NR4 7UH, Norwich, UK; Department of Molecular Microbiology, John Innes Centre, Norwich Research Park, Norwich, NR4 7UH, UK

**Keywords:** *Klebsiella oxytoca*, symbiont, paratransgenesis, 16S rDNA sequencing

## Abstract

Symbioses between bacteria and their insect hosts can range from very loose associations through to obligate interdependence. While fundamental evolutionary insights have been gained from the in-depth study of obligate mutualisms, there is increasing interest in the evolutionary potential of flexible symbiotic associations between hosts and their gut microbiomes. Understanding relationships between microbes and hosts also offers the potential for exploitation for insect control. Here, we investigate the gut microbiome of a global agricultural pest, the Mediterranean fruitfly (*Ceratitis capitata*). We used 16S rRNA profiling to compare the gut microbiomes of laboratory and wild strains raised on different diets and from flies collected from various natural plant hosts. The results showed that medfly guts harbour a fairly simple microbiome, primarily determined by the larval diet in both wild and laboratory flies. However, regardless of the laboratory diet or natural plant host on which flies were raised, *Klebsiella* spp dominated the medfly microbiomes and resisted removal by antibiotic treatment. We sequenced the genome of the dominant putative *Klebsiella* spp (designated ‘Medkleb’) isolated from the gut of the Toliman wild type fruitfly strain. Genome-wide ANI analysis placed Medkleb within the *K. oxytoca / michiganensis* group. Molecular, sequence and phenotypic analyses supported its identity as *K. oxytoca*. Medkleb has a genome size (5 825 435 bp) which is 1.6 standard deviations smaller than the mean genome size of free-living *Klebsiella* spp, and lacks some genes involved in environmental sensing. Moreover, the Medkleb genome contains at least two recently acquired unique genomic islands as well as genes that encode pectinolytic enzymes capable of degrading plant cell walls. This may be advantageous given that the medfly diet includes unripe fruits containing high proportions of pectin. These results suggest that the medfly harbours a commensal gut bacterium that may have developed a mutualistic association with its host and provide nutritional benefits.

## Introduction

All eukaryotic organisms host bacteria (McFall-Ngai et al 2013) and some of the best-studied associations are those that occur between insects and bacteria (Moran et al 2008; McCutcheon and Moran 2011; Moran and Bennett 2014). The majority of microbe-insect interactions are temporary associations. However, some bacteria and insects retain persistent associations over evolutionary time and have evolved co-dependence (Moran et al 2008). In the most extreme examples, these associations have persisted for millions of years and represent intimate co-evolutionary relationships (Moran et al 2005; Moran et al 2008).

Heritable symbioses are one such example and are defined by the direct passage of bacteria from insects to progeny, usually via maternal transmission (Moran et al 2008). These symbioses can be facultative or obligate (Moran et al 2008). Over evolutionary time, facultative symbionts may lose genes that facilitate life in varied environments (Moran and Bennett 2014). A transition to evolutionary interdependence with hosts can often be identified by reduced genome size and reduced GC content (Shigenobu et al 2000; van Ham et al 2003; Wu et al 2006; Hansen et al 2011; Bennett and Moran 2013; Husnik et al 2013). Novel loci can also be acquired via lateral gene transfer or loci deleted through genome reduction (Ochman et al 2000; Ochman and Davalos 2006). Mutualistic benefits provided to insects by bacteria include synthesis of nutrients (Sabree et al 2013; Storelli et al 2018; Sinotte et al 2018), carotenoids (Sloan and Moran 2012), and antipredation molecules (Oliver et al 2003; Sinotte et al 2018). Heritable symbionts include “reproductive parasites” that enhance their fitness by distorting the sex ratios of the hosts’ offspring (Hurst and Frost 2015). For example, male *Zyginidia pullulan* leafhopper embryos are feminised by maternally-inherited *Wolbachia* bacteria (Negri et al 2009). Feminisation benefits *Wolbachia*, because these bacteria preferentially reside within ovaries and therefore are transmitted across generations at higher frequency (Negri et al 2009). The *Buchnera* symbionts of aphids have been extensively studied and represent an example of an advanced obligate mutualism (Shigenobu et al 2000; Tamas et al 2002; van Ham et al 2003). *Buchnera spp* have been associated with aphids for approximately 100M years (Von Dohlen and Moran 2000) and provide the host with essential amino acids (Shigenobu et al 2000). There is an increasing body of research into the identities and potential roles of diverse symbionts found across many different taxa (de Souza et al 2009; Weiss and Kaltenpoth 2016; Holmes et al 2016; Whitten and Dyson 2017; Ballinger and Perlman 2017; Anbutsu et al 2017; Heine et al 2018) including those that are only loosely associated with their hosts, such as the gut symbionts of fruitflies that are the focus of this study.

There are many examples of studies in which the culturable and non-culturable members of gut microbiome communities have been characterised via sequencing of specific regions of the bacterial 16S rDNA. These works show that dipteran gut microbiomes are often relatively stable and simple (Chandler et al 2011; Gould et al 2018; Deguenon et al 2019) and can in some species be strongly influenced by host diet (Chandler et al 2011; Woruba et al 2019). For example, the gut microbiome of *Drosophila melanogaster* is consistently formed by a core of five species: *Lactobacillus plantarum, Lactobacillus brevis, Acetobacter pasteurianus, Acetobacter tropicalis and Acetobacter orientalis* (Gould et al 2018). These simple microbiomes appear to contain bacteria that reliably colonise the gut, suggesting the potential for hosts to actively regulate their gut microbiomes (Lhocine et al 2008; Bosco-Drayon et al 2012; Lindberg et al 2018) and gain potential benefits from doing so (Gould et al 2018). The five core members of the *D. melanogaster* gut microbiome metabolise lactic acid and acetic acid, which may benefit larvae feeding on rotten fruit. In contrast, the *Tephritidae* family of “true fruit-fly” pests hatch in unripe fruit and the culturable species within their gut microbiomes are reported to contain pectinolytic bacteria, which could assist the host in breaking down plant cell walls (Behar et al 2008b; Ben-Yosef et al 2014; Liu et al 2016).

Manipulation of the co-evolved intimacy of symbionts and hosts via paratransgenesis has the potential to be used as a novel mode of pest insect control (Durvasula et al 1997; Whitten et al 2016). Gut symbionts have the potential to augment strategies for control as probiotics. For example, they could boost fitness in insects such as Mediterranean fruitflies (medfly, *Ceratitis capitata*) that are subjected to potentially damaging irradiation as part of sterile insect release control programmes (Jurkevitch 2011). The potential utility of symbionts to either provide new routes for pest control or to improve existing technologies (Leftwich et al 2016) has led to increasing interest in investigating the symbiotic gut microbial communities of key global pests, such as the medfly (Behar et al 2008a; Behar et al 2008b; Ben Ami et al 2010; Gavriel et al 2011), which is the focus of investigation here.

Investigations of culturable gut bacteria in medfly have been predominantly performed using amplified rDNA restriction analysis (ARDRA). These show that *Klebsiella* spp comprise at least 20-30% of the larval, pupal and adult medfly gut microbiomes (Behar et al 2008b; Ben Ami et al 2010; Aharon et al 2013). This has led to the hypothesis that *K. oxytoca* might benefit larval nutrition via its reported pectinolytic activity against fruit sugars, or due to its ability to fix nitrogen (Behar et al 2008b). 16S rDNA analysis has also been used to show that irradiation can alter the microbiome, and particularly diminishes the relative contribution of *Klebsiella* spp (Ben Ami et al 2010). Subsequent reintroduction of *K. oxytoca* to irradiated flies significantly reduced mating latency in comparison to males fed sterile diet (Ben Ami et al 2010), suggesting a potential host fitness benefit.

Although the culturable species within medfly microbiomes have been described, with *Klebsiella* spp appearing to be a typical component, many details remain unknown. For example, we do not yet know the contribution of non-culturable species to both laboratory and wild medfly, whether microbiomes are stable, whether *Klebsiella* spp have the capability to confer a direct fitness benefit to the medfly, and the extent to which *Klebsiella* is heritable. Behar et al. (Behar et al 2008a) suggest that a gut symbiont identified as *K. oxytoca* is heritable and can be transmitted during oviposition. However, *K. oxytoca* was detected in only 1 of 4 replicate samples (Behar et al 2008a) and the potential transmission of *K. oxytoca* during oviposition could not be ruled out . Recovery of GFP labelled *Klebsiella* bacteria in the guts of offspring of mothers into which those bacteria were experimentally introduced provides stronger evidence of vertical transmission (Lauzon et al 2009), though the relative importance of this mechanism is not yet clear. In terms of fitness benefits, Gavriel et al. (Gavriel et al 2011) experimentally depleted the microbiome of male medflies by using irradiation andfed males with a diet either containing *K. oxytoca* (*pro*) or a sterile diet (*ster*). *pro* outcompeted *ster* flies for matings, and females mated to *pro* males were less inclined to re-mate. These data suggest that *Klebsiella* spp could confer host benefits (Behar et al 2008b; Ben Ami et al 2010; Gavriel et al 2011), though it is not yet clear whether this occurs in the natural context.

Here we compared the culturable and non-culturable gut microbiomes of wild-collected adult medflies from a range of different wild hosts, with those of the Toliman wild type laboratory strain reared on a range of different larval diets in the presence and absence of antibiotics. We conducted long-read genome sequencing and analysis of the genome of the dominant, putative *K. oxytoca* (hereafter “Medkleb”) spp extracted from the adult gut of Toliman wild type individuals. This was done to confirm the phylogenetic placement of Medkleb and to interrogate the genome for features characteristic of a nascent evolutionary interdependence with its medfly host. We assembled and annotated the Medkleb genome sequence and tested for signatures of facultative transition, i.e. a reduction in genome size or GC content (Moran et al 2008; McCutcheon and Moran 2011; Moran and Bennett 2014). We conducted comparisons between Medkleb and other *Klebsiella* spp to reveal phenotypes that might potentially facilitate a mutualistic relationship, or indicate restrictions to the environments in which Medkleb might live. We investigated tests of phenotypic features by testing whether Medkleb had the capacity to synthesise secondary metabolites, and by conducting direct biochemical tests for pectinolytic activity.

## Materials and Methods

### 1. Toliman Wild type strain

To directly test the effect of different dietary carbohydrates on the gut microbiome, we raised three replicate samples each of individuals from the Toliman wild type strain on different larval diets. This strain originated from Guatemala and has been reared in the laboratory since 1990. Our Toliman colony has been maintained in non-overlapping generations in a controlled environment room (humidity 50± 5%, temperature 25± 0.5°C) on a 12:12 light:dark cycle for over 30 generations (Leftwich et al 2017). Under this regime, larvae are raised on a sugar-yeast-maize medium (1% agar, 7.4% sugar, 6.7% maize, 4.75% yeast, 2.5% Nipagin (10% in ethanol), 0.2% propionic acid) and adults are given *ad libitum* access to sucrose-yeast food; 3:1 w/w sugar/ yeast hydrolysate, and water.

### 2. Generation of laboratory and wild-derived gut microbiome samples

#### (i) Effect of the larval diet on the adult microbiome in the Toliman wild type

To generate the samples of wild type flies raised on different diets (± antibiotics) for subsequent gut dissection and 16S rRNA amplicon sequencing of the gut microbiome, Toliman flies were cultured from eggs collected over a 24h period placed in one of four larval diet treatments. Three of these diets provided varying carbohydrate levels and sources, while maintaining the same yeast (∼protein) level: 1) Sucrose High Protein (SHP), 2) Glucose & 3) Starch. The fourth diet had a sucrose carbohydrate base but only 60% of the yeast content: Sucrose Low Protein, SLP (Table S1). We included Propionic acid as a food preservative (Leftwich et al 2018). All diet manipulations were done in the presence and absence of antibiotics. For the antibiotic treatments, each larval diet contained a final concentration of 100μg/ml kanamycin, 200 μg/ml ampicillin, 200 μg/ml streptomycin, 50 μg/ml chloramphenicol, 100μg/ml apramycin, 100 μg/ml hygromycin and 200 μg/ml tetracycline. Approximately 500 eggs were placed on 100 mL of the appropriate diet in a glass bottle. When third instar larvae started to “jump” from the larval medium, the bottles were laid horizontally on sand and pupae allowed to emerge for seven days. Pupae were then sieved from the sand and held in 9 mm petri dishes until adult eclosion.

#### (ii) Effect of wild larval diets on the adult microbiome in wild flies under natural conditions

Wild flies were collected at adult eclosion from fallen argan fruit in Arzou, Ait Melloul, Morocco, in July 2014, from Apricots, Oranges and Grapefruits in Chania, Crete, July-September 2014 and from Peaches, Oranges and Tangerines in Ano Lechonia, Greece, July-September 2014. All samples were preserved in 96% ethanol and sent to the UK before gut dissection and DNA extraction for the 16S rRNA amplicon sequencing described below.

### 3. 16S rRNA gene sequencing and bioinformatics analysis of the gut bacteria derived from laboratory- and wild- and derived adult medflies

We analysed the composition of the gut microbiomes in the dissected guts of the laboratory and wild-derived flies describe above, by using 16S rRNA amplicon sequencing. Each of the three biological replicates was a pool of five adult flies (Supplementary Information). Batches of flies were surface sterilized for 30 seconds in 0.5% sodium hypochlorite (bleach) (Sigma-Aldrich, Cat. No.7681529) and washed for 30 seconds in sterile 1M PBS (pH 7.4) three times before being homogenized. 100 µL of the third washes were used to check the surface sterilisation efficiency. There was no microbial growth in any of these tests.

We used sterilised pestles to homogenise the surface-sterilised samples inside 2-mL microcentrifuge tubes, with three freeze/thaw cycles in liquid nitrogen. DNA was extracted using the DNeasy Blood and Tissue Kit (Qiagen) and quality checked using a NanoDrop (Thermoscientific Nanodrop 8000 Spectrophotometer). Approximately 100 ng of DNA per sample was used as the template for PCR amplification with bacterial universal primers 515F (5′-GTG CCA GCM GCC GCG GTA A-3′) and 806R (5′-GGA CTA CHV GGG TWT CTA AT-3) against the 16S rRNA gene. Amplicon sequencing was performed using paired-end 250 bp V2 chemistry (Illumina MiSeq platform, Earlham Institute provider).

Demultiplexed sequences were obtained using mothur v38.2 (Schloss et al 2009), following their standard MiSeq operating procedures. Sequence variants were assigned to operational taxonomic units (OTUs) at a 97% similarity threshold. Taxonomy assignment of OTUs using the Silva database (release 132). The minimum library sizes per sample were ∼17K after passing quality control. All statistical analyses of amplicon data were conducted in R v3.6.2 (R Core Team 2019) using the phyloseq (McMurdie and Holmes 2013), vegan (Oksanen et al 2007) and tidyverse (Wickham 2017) packages. Sequences were rarefied to normalise library sizes. Alpha diversity was estimated using the Shannon species diversity index. We visualised differences in bacterial community structure among samples (beta diversity) using non-metric multidimensional scaling (NMDS) plots of Bray-Curtis distances and performed multivariate analysis of variance (PERMANOVA) with 999 permutations on Unifrac distance matrices.

### 4. Genome sequencing of the dominant gut microbiome *Klebsiella* spp symbiont (Medkleb) from the Toliman wild type

We investigated the identity of the recurrent *Klebsiella* spp bacterial symbiont through isolation of culturable *Klebsiella* spp. colonies from the Toliman strain.

#### (i) Clonal isolation

Clonal isolates of *Klebsiella* spp obtained from individuals of the Toliman wild type strain were made by taking surface sterilized, homogenised samples from adults reared under the standard conditions described above and plating them onto Simmon’s Citrate LB Agar (with bromothymol blue as a colour indicator). This is a substrate recommended for the isolation of *Klebsiella oxytoca and K. pneumoniae* (Simmons 1926). Culture plates were made from 15 biological replicates of medfly. Cultures were checked for morphological uniformity and their identity confirmed with PCR amplification with universal bacterial primers 28F and 806R. Thirteen of the 15 isolates had identical 16S rRNA gene sequences and were BLAST matched to *K. oxytoca* and our most abundant OTU from 16S rRNA amplicon sequencing. Two isolates contained colonies which BLAST matched to *Pantoea* spp. We chose a single *Klebsiella* spp colony at random for genomic sequencing, as described below.

#### (ii) DNA preparation

The single clonal isolate selected for genome sequencing was streaked onto LB media (15g/L Agar; 5g/L NaCl; 5g/L yeast extract; 1.5g/L glucose; 10g/L tryptone) and incubated overnight at 25°C, then transferred into a 1.5ml microcentrifuge tube containing 1ml of 10% glycerol. The sample was vortexed for 30 secs then centrifuged at 12000 RPM for 10min. Glycerol was then removed, and the pelleted bacteria re-suspended in 2ml of SET buffer (65% v/v molecular grade H_2_ O (ThermoFisher); 20% v/v Tris (pH8); 5% v/v 5M NaCl; 5% v/v 10% SDS; 5% v/v 0.5M EDTA) before transfer to a 15ml falcon tube. 20μg of lysozyme (Sigma) and 0.4μg achromopeptidase (Sigma) suspended in 40µl of molecular grade water (ThermoFisher) and 0.02μg of RNase (Fermentas) were added. The sample was mixed gently then incubated at 37°C for 2hrs. 240µL of 10% sodium dodecyl sulphate and 56µL of proteinase K (20mg/ml) were added, before a second incubation at 56°C for 2hrs, with manual mixing every 30 mins. 800µL of 5M NaCl and 2ml of chloroform were added and the sample was mixed by hand for 10 mins, before centrifugation at 4000 RPM for 12 mins. The aqueous phase was then carefully transferred to a fresh tube. DNA was precipitated in 0.6 volume isopropanol, then transferred to a 1.5ml microfuge tube by pipette. DNA was washed once with 70% ethanol. Ethanol was removed, then 1ml of 70% ethanol was added, and DNA was left to incubate overnight at 4°C. Ethanol was again removed, and DNA was re-suspended in 200µL of molecular grade water (ThermoFisher).

#### (iii) Single molecule real time (SMRT/PacBio) Medkleb genome sequencing

DNA purity, concentration, and average fragment size were analysed using Nanodrop (ThermoFisher), Qubit v2.0 (Invitrogen) and Agilent Tapestation 4200 (Agilent) respectively. DNA was fragmented using a G-tube (Covaris), and SMRTbell library construction was carried out using a Template Prep Kit 1.0 (PacBio). The library was then size selected to >7kb using the BluePippin system (Sage Science). Sequencing was carried out on a Pacific Biosciences RSII instrument, using two RSII SMRTcells v3 and P6-C4 chemistry (PacBio, Earlham Institute provider). Each cell was sequenced using a 240-minute movie, using the Magbead OCPW v1 protocol (PacBio).

#### (iv) Medkleb genome assembly

The Medkleb genome was assembled according to the Hierarchical Genome-Assembly Process (HGAP.3) protocol (Chin et al. 2013) as follows. 1) Mapping – BLASR (Chaisson et al., 2012) was used to map reads >500bp with a read quality >0.8 to seed reads >6000bp. 2) Pre-assembly -the Directed Acyclic Graph Consensus (DAGCon) algorithm (Lee et al 2002) was used to produce a consensus sequence based on BLASR mapping. DAGCon then trimmed the consensus, producing an error-corrected pre-assembled read. 3) *de novo* genome assembly -the overlap-layout-consensus assembler Celera Assembler v8.1 was used to process the pre-assembled read into a draft assembly. 4) Final consensus -the draft assembly was polished using the Quiver multiread consensus algorithm (Chin et al 2013). 5) The final consensus sequence was then manually trimmed to circularise the genome and place the stop codon (TGA) of the *dnaA* gene at the 5’ terminus.

#### (v) Medkleb genome quality control

An estimated Quiver quality value (QV) for the Medkleb final consensus genome was provided (Earlham Institute). Genome completeness was estimated with both benchmarking universal single copy orthologues (BUSCO) software 3.0.0 (Simão et al 2015), and CheckM (Parks et al 2015). The *Enterobacteriales* order and *Enterobacteriaceae* family were used as reference datasets for BUSCO and CheckM analyses respectively. Genome contamination was estimated with CheckM, and mlplasmids (Arredondo-Alonso et al 2018) was used to classify contigs as either chromosomal or plasmid DNA. The *Klebsiella pneumoniae* support-vector machine (SVM) model was utilised for the analysis, with minimum posterior probability specified at 0.7 and minimum contig length at 1,000nt.

#### (vi) Annotation and genome mapping

Coding sequences within the Medkleb chromosomal DNA and plasmids, were called with the Prodigal algorithm (Hyatt et al 2010). Gene calls were then annotated with Classic-RAST (Overbeek et al 2014). Ribosomal RNAs (rRNAs) and transfer RNAs (tRNAs) were called and annotated with Classic-RAST. Circular maps were created for the Medkleb chromosomal DNA and plasmids using DNAplotter (Carver et al 2009).

### 5. Phylogenetic analyses of the Medkleb *Klebsiella* gut symbiont genome sequence

#### (i) 16s rRNA gene sequence analyses

The Medkleb genome was searched for regions homologous to the 16S rRNA gene sequence of *K. oxytoca* strain ATCC 13182 (NR_118853.1) with BLASTn (Altschul et al 1997). The region with greatest homology to NR_118853.1 (nts 341370-342821) was then parsed with RNAmmer 1.2 (Lagesen et al 2007) which predicts ribosomal genes. Sixty-one 16S rRNA gene nucleotide sequences representing 60 *Klebsiella* strains, *Pseudomonas aeruginosa* strain JB2 and a putative Medkleb 16S sequence were used to create a phylogeny with the SILVA ACT web app (Pruesse et al 2012). Where possible, non-redundant sequences were extracted from the SILVA rRNA gene database (Quast et al 2013). All sequences were almost complete (>1400bp) and met the standard operating procedure for phylogenetic inference (SOPPI) quality criteria set out by Peplies et al. (Peplies et al 2008). The tree was computed with the FastTree2 maximum likelihood programme (Price et al 2010) using the GTR evolutionary model and gamma distribution parameters (Yang 1994). The resulting phylogeny was constructed with FigTree 1.4.3 (Rambaut and Drummond 2009).

#### (ii) Average Nucleotide Identity (ANI) analyses

35 RefSeq whole genome entries extracted from the NCBI database (Geer et al 2010), representing four *Klebsiella* species, were used to calculate a hierarchical clustering based on Average Nucleotide Identity (Konstantinidis and Tiedje 2005). The ANI Calculator (Figueras et al 2014) was used to compute the hierarchy using the BIONJ algorithm (Gascuel 1997). The tree was constructed with FigTree 1.4.3 (Rambaut and Drummond 2009). To predict recently acquired Medkleb sequences, the genome was aligned with three closely related bacteria. Genomes were manually re-ordered to place the *dnaA* stop codon at the 5’ terminus. Synteny was then predicted with progressive Mauve (Darling et al 2010).

### 6. Metabolic functions of the Medkleb *Klebsiella* gut symbiont

#### (i) Analysis of pectinolytic enzyme activity

Medkleb’s ability to degrade pectin was compared to two bacterial species identified as positive and negative controls; *Erwinia carotovora* (+ve control) and *Rhizobium leguminosarum* (-ve control) (Wegener 2002; Xie et al 2012). Medkleb bacteria were cultured in LB broth (5g/L NaCl; 5g/L yeast extract; 1.5g/L glucose; 10g/L tryptone) which had been stored in 50ml glass bottles and autoclaved prior to use. Each bottle was inoculated with a “loop” of bacteria and incubated, shaking, in an orbital incubator (New Brunswick Scientific Innova 44) at 200RPM. Medkleb and *Erwinia carotovora* were incubated at 37°C and *Rhizobium leguminosarum* was incubated at 28°C until optical density was greater than 1.0 at 600nm.

#### (ii) Analysis of presence of pehX and 16S rRNA genes in medkleb

Bacterial DNA was extracted using a DNeasy blood and tissue kit (Qiagen) and microbe lysis buffer (MLB) (20 mg/ml of lysozyme (Sigma) and 5mg/ml of achromopeptidase (Sigma) in 20 mM Tris-HCl, 2 mM EDTA, 1.2% Triton X (pH 8.0)). 2ml of Medkleb, *Erwinia carotovora* and *Rhizobium leguminosarum* cultures were centrifuged at 13K RPM for 5 mins, before the supernatants were removed. Pellets were then homogenised with a clean pestle in liquid nitrogen. 180μl of MLB was added before the sample was vortexed and incubated at 37°C for two hours. Samples were vortexed every 30 mins during the two-hour incubation. Buffer AL and ethanol were mixed 1:1 (Buffer ALE) and warmed to 55°C. Each sample had 400μl of warm Buffer ALE added before being immediately vortexed for 10-15 secs. Samples were transferred to a spin column and centrifuged at 8000RPM for 60 secs. 500μl of Buffer AW1 was added and the sample was centrifuged again at 8000RPM for 60 secs. 500μl of Buffer AW2 was added and the samples were centrifuged at 13 000 RPM for 4 mins. 35μl of warm Buffer AE (60°C) was added to the centre of the spin membrane and the samples were centrifuged at 6000RPM for 60 secs. DNA purity and concentration were measured using a Nanodrop (ThermoFisher). The *K. oxytoca* polygalacturonase gene pehX (AY065648.1) was aligned to genomes of *Klebsiella* bacteria using BLAST (Altschul et al 1997). The presence of polygalacturonases in genomes of *Klebsiella* bacteria was assessed using the Carbohydrate-Active enZymes database (CAZy) (Lombard et al 2010).

#### (iii) Polygalacturonase enzyme assay

Polygalacturonase production of Medkleb, *Erwinia carotovora* and *Rhizobium leguminosarum* was measured using a DNS colorimetric method (Miller 1959) with a protocol adapted from Sohail et al. (Sohail and Latif 2016) and Sigma Aldrich protocol EC 3.2.1.1. Cultures were diluted with LB broth to an optical density of 1.0 at 600nm. Bacteria were then filtered from culture media with 0.2μm PES syringe filters (ThermoFisher). Treatment reactions (which were run in triplicate) were set up with 1ml of appropriate filtrate and 1ml of polysaccharide solution (PS) (0.9% polygalacturonic acid (ThermoFisher) in 0.1M sodium acetate (ThermoFisher) (pH 4.5)), a blank reaction was set up with 1ml of PS only. All reactions were incubated at 45°C for 30mins before 1ml of colour reagent solution (20% 5.3M potassium sodium tartrate, tetrahydrate in 2M sodium hydroxide solution; 50% 96 mM 3,5-Dinitrosalicylic acid solution; 30% molecular water (ThermoFisher)) was added. All reactions were incubated at 100°C for 15 mins, then placed on ice to cool to room temperature. Once cooled, 12ml of molecular water (ThermoFisher) was added to each reaction, followed by hand mixing. Absorbance at 530nm (ΔA530) was measured using a spectrophotometer (Biochrom) which had been blanked for air, with the corrected ΔA530 for treatment reactions being calculated as:

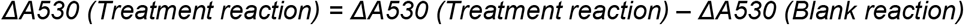

Units of polygalacturonase in the filtrate were calculated via comparison to a standard curve of galacturonic acid (Sigma). Standards were made with between 50μl and 1 ml of monosaccharide solution (MS) (1.8% galacturonic acid (ThermoFisher) in 0.1M sodium acetate (ThermoFisher) (pH 4.5) and topped up to 2ml total volume with molecular water (ThermoFisher). A standard blank was set up containing 2ml of molecular biology grade water only. 1ml of colour reagent solution was then added before incubation at 100°C for 15 mins. Standards were placed on ice to cool to room temperature before 12ml of molecular water was added and ΔA530 was measured using a spectrophotometer (Biochrom) which had been blanked for air. The corrected ΔA530 for standards was calculated as: *ΔA530 (Standard) = A530 (Standard) – A530 (Blank)*. The standard curve was used to estimate mg of galactose released in treatment reactions with linear regression, and units of polygalacturonase per ml of filtrate were then calculated as: *Units/ml enzyme = (mg of galactose released)/ml of filtrate)*.

#### (iv) Prediction of higher order metabolic functions and secondary metabolites

The higher order metabolic functions of genes were predicted using the Kyoto Encyclopaedia of Genes and Genomes (KEGG) (Kanehisa et al 2016). Secondary metabolites were predicted using antiSMASH 6.0 beta (Blin et al 2019).

## Results

### 1. Characterisation of the medfly gut microbiome using amplicon sequencing

#### (i) Bacterial community diversity

We found that OTU richness and diversity varied significantly among host samples according to treatment and origin (Figure 1 A &B). Community structure (beta diversity) was affected primarily by diet, in both wild and lab flies (PERMANOVA: F_8,48_ = 2.74, P <0.001, R^2^ = 0.27, Figure 1A). Within lab flies, age (F_1,30_ = 4.9, P<0.001, R^2^ = 0.18), diet (F_3,30_ = 2.03, P =0.023, R^2^ = 0.12), and antibiotic treatment (F_1,30_ = 4.9, P =0.002, R^2^ = 0.09), all affected community structure, with decreasing levels of effect. Bacterial species diversity (alpha diversity) did not vary significantly in the wild or laboratory population strains but was lower in antibiotic treated flies. However, the Shannon index was not significantly different between any samples (Figure 1B). Overall, we found no evidence of large-scale changes in diversity or composition of the microbiome despite testing multiple wild food sources, effects of laboratory diets and antibiotic treatment (Figure 1C).

**Figure 1.**
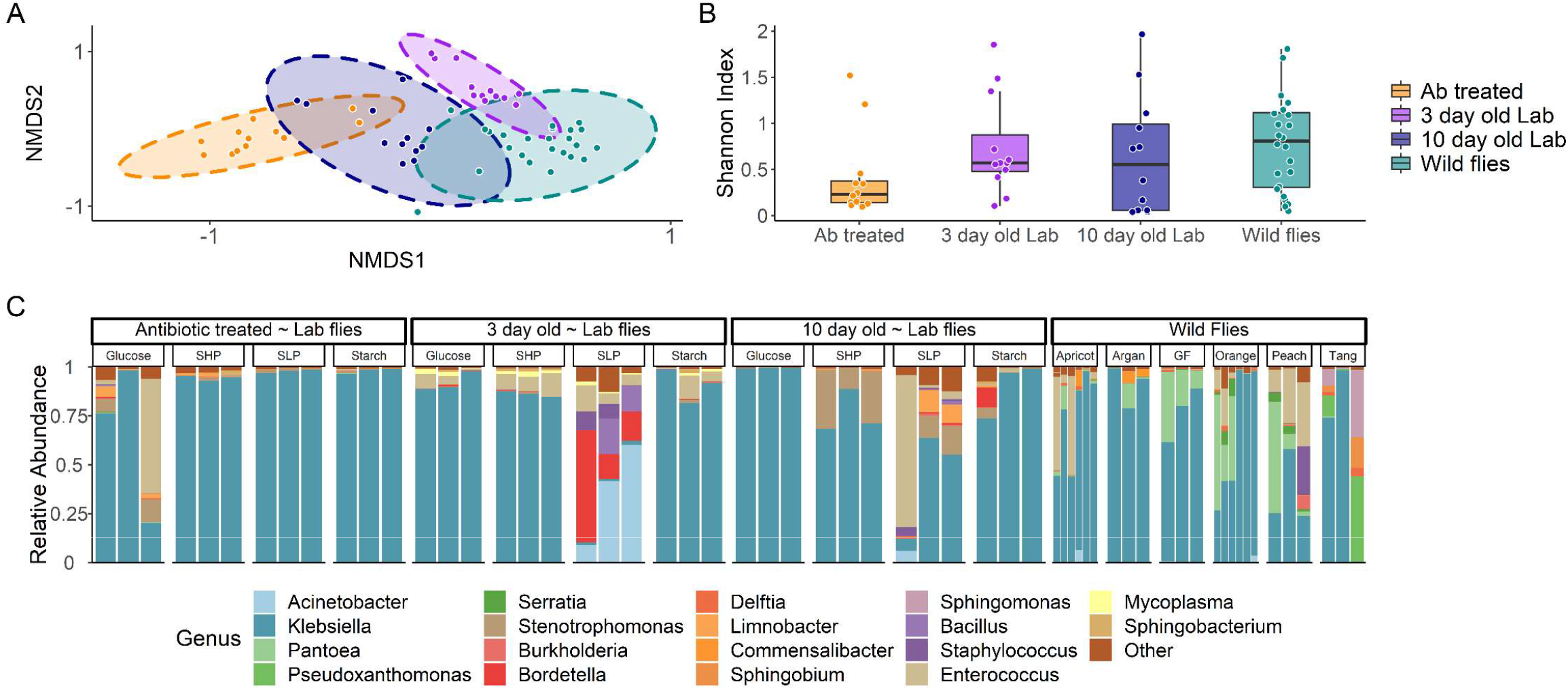
16S rRNA profiles of wild and laboratory strains of medfly on varied larval diets and in the presence and absence of antibiotic treatment. Microbiome composition was measured as **(A)** community structure/beta diversity visualised as NMDS plots using a Bray-Curtis Dissimilarity Index with 95% confidence ellipses, **(B)** species richness/alpha diversity using the Shannon Index. Boxplot displays median, hinges are first and third quartiles, whiskers extend from hinge to 1.5* the interquartile range. **(C)** Bar plot of microbiome profiles. Ab = antibiotic treated (100μg/ml kanamycin, 200 μg/ml ampicillin, 200 μg/ml streptomycin, 50 μg/ml chloramphenicol, 100μg/ml apramycin, 100 μg/ml hygromycin and 200 μg/ml tetracycline), 3 day old Lab = 3 day old adults from the Toliman laboratory strain; 10 day old Lab = 10 day old adults from the Toliman laboratory strain, wild flies = flies from fallen argan fruit in Arzou, Ait Melloul, Morocco; Apricots, Oranges and Grapefruits in Chania, Crete; and from Peaches, Oranges and Tangerines in Ano Lechonia, Greece (n = 5 flies per replicate, 3 biological replicates).

#### (ii) Dominant bacterial taxa

Four bacterial families representing two bacterial phyla made up over 90% of the sequences in our dataset. These were the Proteobacteria, Enterobacteriaceae (79%), Moraxellaceae (2.3%) and Xanthomonadaceae (3.6%), and the Firmicutes, Enterococcus (7.3%) (Figure 1C). Of these, a single bacterial genus *Klebsiella* spp emerged as a core member of the bacterial microbiome. A putative *Klebsiella* spp was found in every medfly population sample and comprised 73.6% of the entire dataset. This suggests that, although medflies are extremely polyphagous, they have a stable microbiome, containing a recurrent *Klebsiella* spp symbiont.

### 2. Medkleb genome sequencing

#### (i) Classification of Medkleb sequencing contigs

Total Medkleb DNA was sequenced on a PacBio RSII module and assembled using the HGAP.3 algorithm and Quiver (Chin et al., 2013). This process detected one large contig that sequestered 94% of total gene space (Figure 2) and four small contigs (mkp1-4; Figure S1). This suggests that the total Medkleb DNA complement is formed of one chromosome and four plasmids. Consistent with this, the mlplasmids software (Arredondo-Alonso et al., 2018) classified the large Medkleb contig as chromosomal and the four smaller contigs as plasmids (Table S2). In addition, mkp2, mkp3 and mkp4 were sequenced with relatively high coverage depth (Figure S2) and showed evidence for high copy numbers, a common plasmid trait (Providenti et al., 2006). Finally, mkp3 and mkp5 were demonstrated to have low GC content relative to the putative chromosome (Table S2), which again is characteristic of plasmid DNA (Nishida 2012). mkp2 and mkp4 both exhibited a similar GC content to that of the chromosome, suggesting that they have been acquired more recently (Rocha and Danchin 2002).

**Figure 2.**
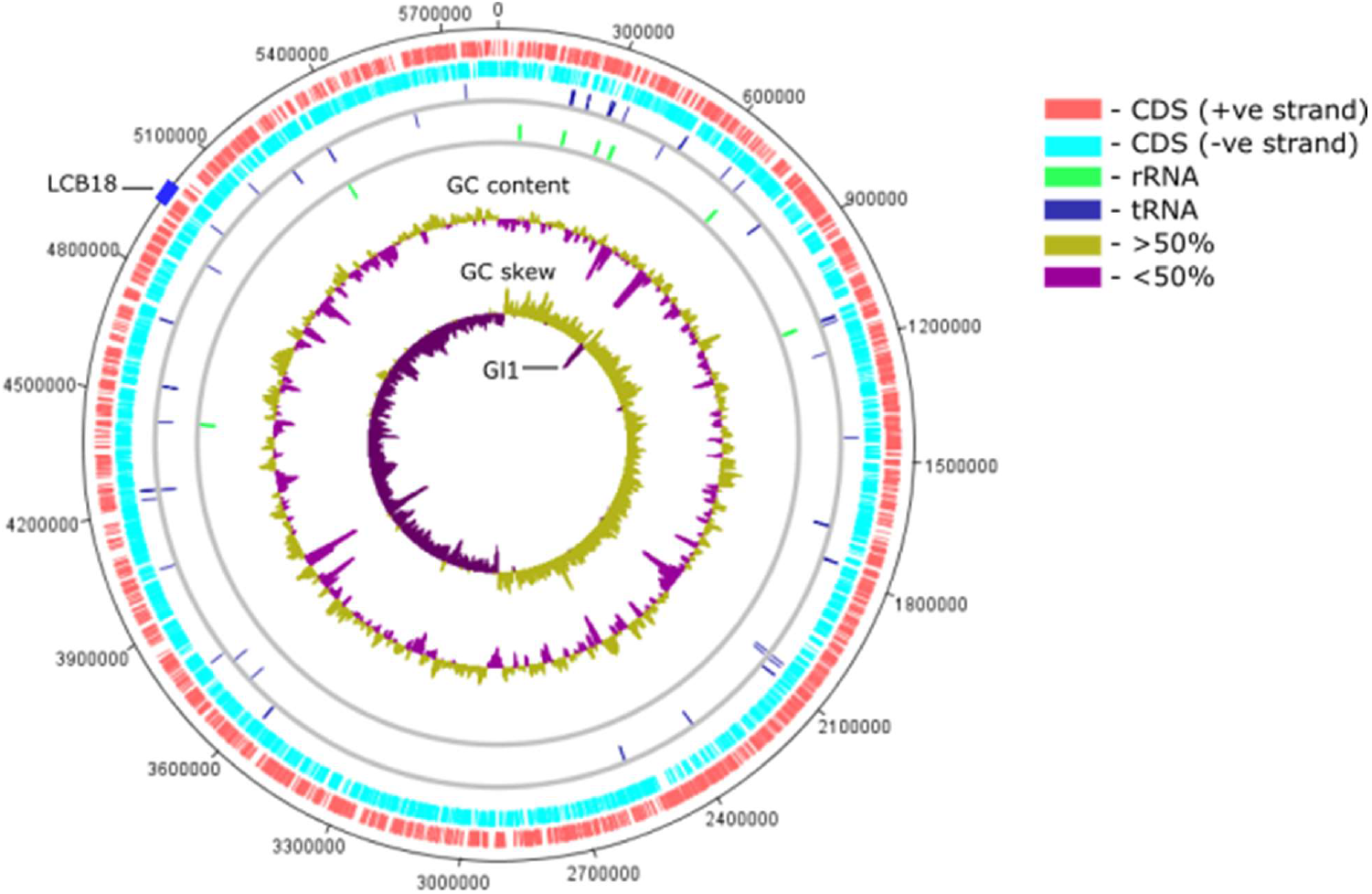
Circular summary map of the Medkleb chromosome. The Medkleb chromosome is represented with the stop codon (TGA) of the *dnaA* gene at position 0. Track one; 2 541 red ticks represent gene coding sequences on the positive strand. Track two; 2847 light blue ticks represent gene coding sequences on the negative strand. Track three; 50 dark blue ticks above the grey line represent tRNAs on the positive strand, and 35 ticks below the line represent tRNAs on the negative strand. Track four; 18 green ticks above the grey line represent rRNAs on the positive strand, and 7 ticks below the line represent rRNAs on the negative strand. 64% of rRNAs are found close to *oriC*. Track five – GC content; regions containing >50% GC content are mustard and regions containing <50% GC content are purple. Track six – GC skew; regions containing >50% Gs are mustard and regions containing <50% Gs are purple. GC skew was asymmetric between leading and lagging strands, with the purple spike between bases 654 265-677 695 (GI1) representing recent horizontal gene transfer (Lawrence and Ochman 1997). The LCB18 gene island (≈nts 4.97 × 10^6^ ∼ 5 × 10^6^) is represented by a blue block on the outer ring.

#### (ii) Quality control

PacBio coverage depth >100X is considered sufficient for resolving nucleotide sequences (Rhoads et al., 2015) and this threshold was met by all contigs other than mkp4 (Figure S2). The QV sequence resolution was 48.9 (an average error rate of 1 base in every 80100) and hence the Medkleb chromosome was of high quality (Figure S2). The plasmid QV’s were: mkp1) 48, mkp2) 47.3, mkp3) 45.3 and mkp4) 44.8 (Figure S2). Although the plasmid sequences had lower resolution, they were robust, with accuracy >99.994% in all cases. The completeness of the Medkleb genome was measured with BUSCO (Simão et al 2015), which was used to search the assembly for 781 marker genes associated with bacteria of the *Enterobacteriales* order. BUSCO estimated that the Medkleb chromosomal sequence was 99.5% complete (777 of 781 genes complete and single copy), far exceeding general quality thresholds (Bowers et al 2017). Furthermore, three genes that were predicted by BUSCO to be fragmented were likely heterozygous alleles that failed to collapse during the annotation. CheckM (Parks et al 2015) was used as a second method to assess the Medkleb chromosome for completeness and contamination. By using as reference 1162 marker genes associated with the family *Enterobacteriaceae*, the CheckM software estimated that the Medkleb genome was 99.7% complete and 0.212% contaminated, again far exceeding standard quality thresholds (Bowers et al 2017).

#### (iii) Medkleb genome

The general characteristics of the Medkleb genome were consistent with those published as *Klebsiella oxytoca* (Shin et al 2012; Bao et al 2013). The Medkleb genome was 5 825 435 nt in length, with 5 388 coding sequences and a GC content of 56.03% (Figure 2; Table S3). At 87.8%, overall coding sequence was within the expected range (Kuo et al 2009), and, as predicted by Reva et al. (2004), genes were distributed symmetrically between the two DNA strands. There were 2 541 coding sequences on the positive strand, which were predicted to code for 2 473 proteins, 50 tRNAs and 18 rRNAs. On the negative strand there were 2 847 coding sequences which coded for 2 805 proteins, 35 tRNAs and 7 rRNAs. KEGG (Kanehisa et al 2016) predicts that the Medkleb genome encodes for genes with 2 842 distinct molecular functions. The genes for 16 (64%) ribosomal RNAs (rRNA) clustered between nts 62 435 – 70 701 near the origin of replication (*oriC*). Medkleb’s GC content was 57.28% for protein coding genes, 53.83% for rRNA and 58.93% for tRNA. Consistent with Lobry (1996) GC skew was asymmetric, with an overrepresentation of Gs on the leading strand and Cs on the lagging strand, indicating that the genome is largely stable with few recent recombination events. However, there is one obvious exception, in which GC skew was inverted (>50% C’s) between bases 654 265-677 695. This is indicative of a recent introgression that has resulted in the acquisition of a new gene island (designated GI1) (Lawrence and Ochman 1997; Wixon 2001).

#### (iv) Medkleb plasmids

The lengths and GC contents of mkps 1-4 (Table S2) were all within expected range for plasmids associated with *Klebsiella oxytoca*. mkps 1-4 are predicted by KEGG (Kanehisa et al 2016) to code for 48 functional orthologues and devote 14-22% of gene space to plasmid associated genes and mobile element coding sequences. This is substantial in comparison to the main chromosome, which only allocated 0.02% to such features. mkps 1, 3 and 4 exhibited asymmetric gene distribution between DNA strands (coding bias), which is common for plasmid genomes (Reva and Tümmler 2004). The coding bias of mkp3 was particularly clear, with ∼90% of the total gene complement found on the positive strand. mkp4 was the only plasmid predicted by antiSMASH 6.0 beta (Blin et al 2019) to code for secondary metabolites, which included cloacin (de Graaf et al 1969) and colicin bacteriocins (Cascales et al 2007).

### 3. Taxonomic placement of the Medkleb *Klebsiella* gut symbiont

#### (i) Taxonomic identification of Medkleb –16S rDNA and ANI analysis

Medkleb was predicted to be a strain of *K. oxytoca*, which has previously been proposed as a major component of the medfly microbiome (Behar et al 2008a). The 16S rRNA gene sequence of *K. oxytoca* strain ATCC 13182 (NR_118853.1) was used to locate homologous sequences in the Medkleb genome using the BLASTn algorithm (Altschul et al 1997). This revealed that the Medkleb genome contains eight sequences >99% related to NR_118853.1, which were all predicted to code for 16S rRNAs by RNAmmer 1.2 (Lagesen et al 2007). Nucleotides 341370-342821 (mk16S) which had the greatest homology with NR_118853.1 were therefore used to represent Medkleb in subsequent taxonomic analyses. These sequences were used in an initial taxonomic description (as described in the Supplementary Information) which indicated that Medkleb was indeed likely to fall within a group of *Klebsiella oxytoca / michiganensis* spp (Figure S3).

A more stringent Average Nucleotide Identity (ANI) (Konstantinidis et al., 2005) analysis was also conducted. The Medkleb genome was positioned in an ANI matrix with 35 RefSeq *Klebsiella* genomes that were extracted from the NCBI database (Geer et al 2010) (Figure 3). This analysis placed Medkleb in a lineage with 13 strains classified as both *K. oxytoca* and *K. michiganensis*. ANI scores >95% are required to classify bacteria as the same species (Richter and Rosselló-Móra 2009; Kim et al 2014). This 95% similarity threshold was met by all thirteen members of Medkleb’s ANI clade (henceforth referred to as the Medkleb group) (Figure S4). Hence both 16S rRNA gene and ANI analyses did not differentiate between *K. oxytoca* and *K. michiganensis* as they are currently named.

**Figure 3.**
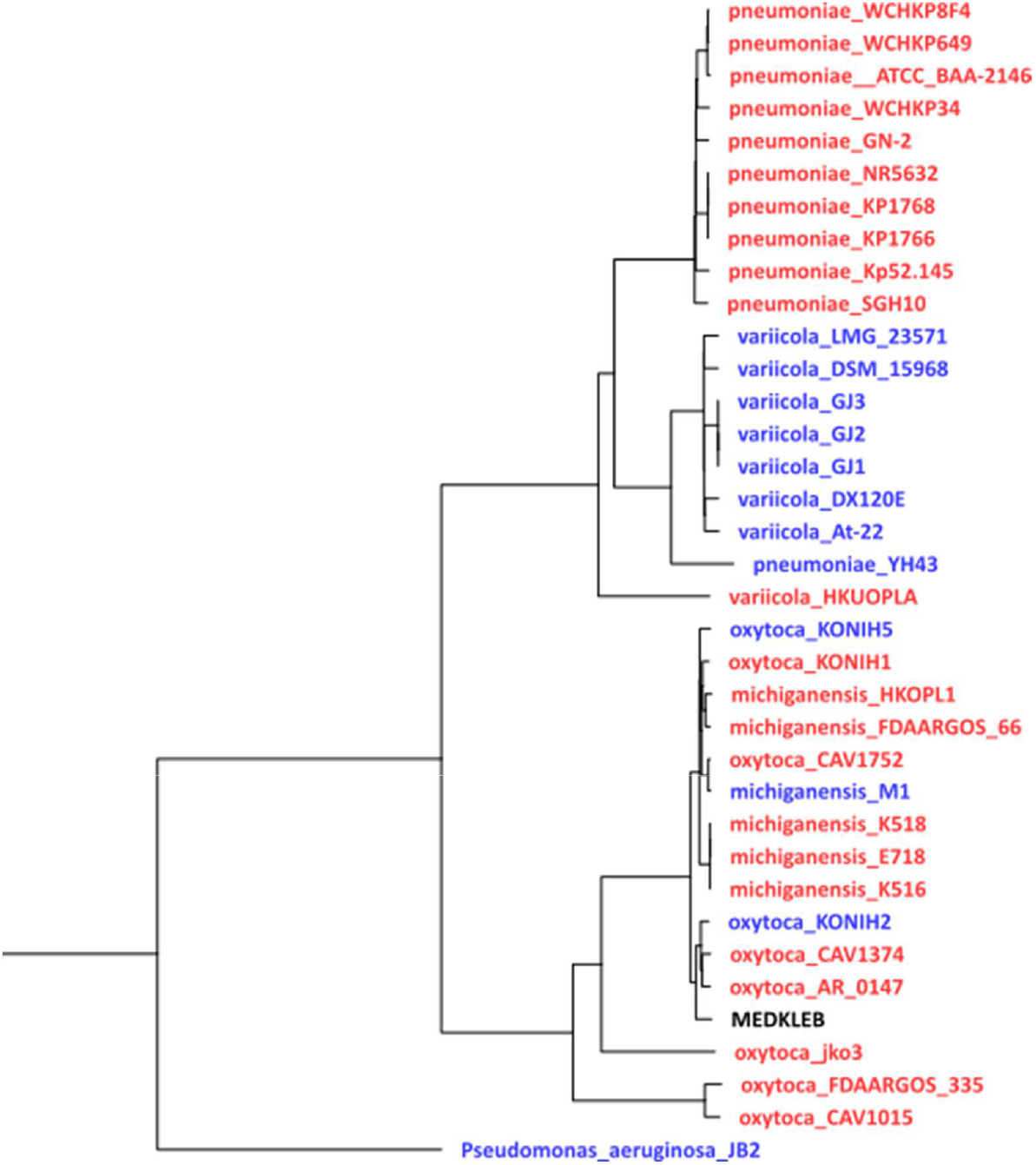
ANI hierarchical clustering showing the evolutionary relationship of environmentally-derived and host-derived *Klebsiella* bacteria. The tree was created using the ANI calculator (Figueras et al 2014), with *Pseudomonas aeruginosa* strain JB2 selected as the outgroup. Bacteria derived from animal hosts (red and black), and environmentally derived bacteria (blue), generally fell into three clades: 1) *K. pneumoniae*, 2) *K. variicola* and 3) *K. oxytoca/michiganensis*. With one exception in each group (YH43 and HKUOPLA), all *K. pneumoniae* are host derived and all *K. variicola* are environmentally derived. *K. oxytoca/K. michiganensis* have been isolated from both environmental and animal sources, but their sequences did not cluster according to source status. According to the ANI species threshold set by Kim et al. (2014), Medkleb is conspecific with twelve strains which have been classified as both *K. oxytoca* and *K. michiganensis*.

### 4. Metabolic functions of the Medkleb *Klebsiella* gut symbiont

#### (i) Bioinformatic analyses of pectic lyases in Medkleb

The ability to degrade pectin is a defining phenotype of *K. oxytoca* and is proposed to be conferred by the polygalacturonase enzyme, pehX (AY065648.1) (Kovtunovych et al 2003)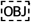We investigated whether *K. oxytoca* and *K. michiganensis strains* of the Medkleb group could be distinguished via homology to pehX but our analysis detected no significant differences between strains (analysed with BLAST; signed-rank linear model F_1,11_ = 2.57, p = 0.137). Unexpectedly, *K. michiganensis* shared greater homology with AY065648.1 than did *K. oxytoca* (Table 1), despite the fact that *K. michiganensis* does not degrade pectin (Saha et al 2013). In addition, the CAZy database (Lombard et al 2010) identified two pectate lyases (one from family 2 and one from family 9) in all Medkleb group genomes available on the database (Table S5).

**Table 1.** Genomic features of the ‘Medkleb group’ of bacteria. *pehX* sequence identity (calculated with BLAST (Altschul et al., 1997)) was not significantly different between species (analysed with BLAST; signed-rank linear model F_1,11_ = 2.57, p = 0.137). *K. michiganensis* genomes contained sequences more closely related to AY065648.1 than was found for *K. oxytoca* genomes (mean relatedness = 88.4% vs 87.9%). All genomes available on the CAZy database (Lombard et al 2010) are predicted to code for 2 pectate lyases. Medkleb had the smallest genome in the group^1^, and the second highest GC content^2^.

| Species | Strain | Genome size (nt) | GC (%) | *pehX* identity (%) | CAZy lyases |
| --- | --- | --- | --- | --- | --- |
| *oxytoca* | Medkleb | 5825435<sup>1</sup> | 56.03 | 87.43 | NA |
| *michiganensis* | M1 | 5865090 | 56.13<sup>2</sup> | 88.47 | 2 |
| *michiganensis* | HKOPL1 | 5914407 | 55.92 | 88.56 | 2 |
| *oxytoca* | CAV1752 | 5992008 | 55.16 | 88.47 | 2 |
| *michiganensis* | FDAARGOS_66 | 6071464 | 55.94 | 88.39 | NA |
| *michiganensis* | E718 | 6097032 | 56.02 | 88.31 | 2 |
| *michiganensis* | K518 | 6138996 | 55.95 | 88.39 | 2 |
| *michiganensis* | K516 | 6139574 | 55.96 | 88.31 | 2 |
| *oxytoca* | KONIH1 | 6152190 | 55.91 | 88.61 | 2 |
| *oxytoca* | KONIH5 | 6179177 | 55.81 | 87.82 | 2 |
| *oxytoca* | KONIH2 | 6190364 | 55.77 | 87.43 | 2 |
| *oxytoca* | CAV1374 | 6257473 | 55.75 | 87.87 | 2 |
| *oxytoca* | AR_0147 | 6350620 | 55.57 | 87.89 | 2 |

<sup>1</sup> smallest genome
<sup>2</sup> highest GC content

#### (ii) Polygalacturonase enzyme assay

Medkleb was pehX positive when analysed biochemically (Figure S6), which suggested that it should be classified as *K. oxytoca* and can degrade pectin (Kovtunovych et al 2003; Saha et al 2013). We quantified Medkleb’s capacity to degrade polygalacturonic acid-in comparison to positive (*E carotovora)* and negative (*R* leguminosarum) control specimens, using a DNS colorimetric method (Miller 1959). A standard curve of galacturonic acid incubated with DNS (Figure 4A), showed a positive relationship with colour change when the OD was measured at 530nm (linear model, F_1,4_ = 176.4, p <0.001, R^2^ = 0.97). Filtrate of Medkleb and *E. carotovora* culture media both contained pectinolytic enzymes, producing measurable colour change when incubated with polygalacturonic acid and DNS (Figure 4B; (Sohail and Latif 2016)). Medkleb filtrate reduced polygalacturonic acid with around 65% the efficiency of the *Erwinia carotovora*, but *R leguminosarum* filtrate did not demonstrate any measurable pectinolytic activity (Figure 4C). These data support the placement of Medkleb as *K. oxytoca*.

**Figure 4.**
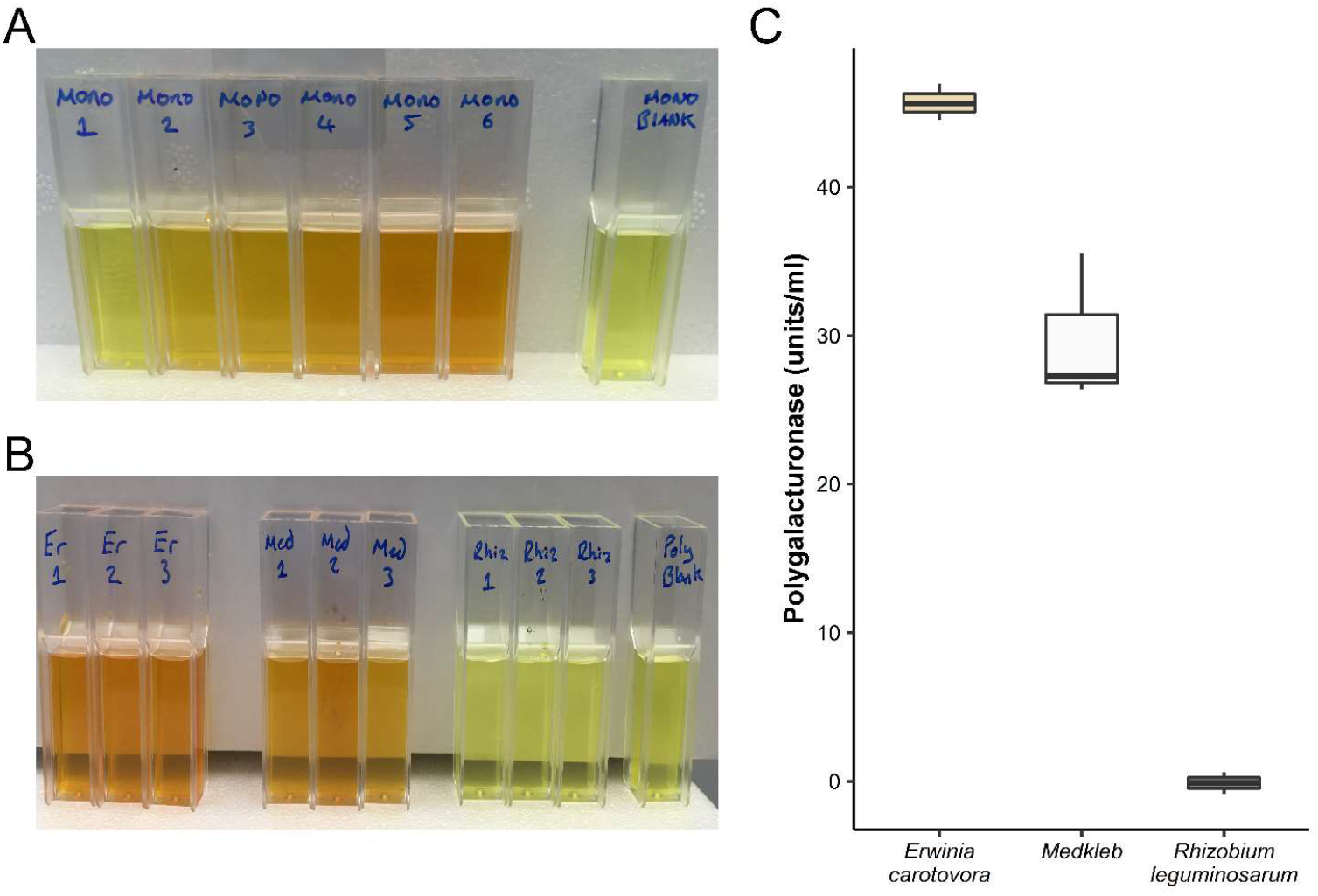
Polygalacturonase production by *Erwinia carotovora* (Er), Medkleb (Med) and *Rhizobium leguminosarum* (Rhiz). A) Monosaccharide colorimetric reaction mixtures, standard curve – The quantity of sugar in standards ranged from 0.9mg (mono 1) to 18mg (mono 6). Mono blank contained no sugar. When incubated with DNS, colour change (measured at ΔA530) occurred for all reactions relative to the blank. The relationship between sugar quantity and colour change was significant (linear model, F_1,4_ = 176.4, p <0.001, R^2^ = 0.98). B) Bacteria reaction mixtures – Polygalacturonase production of three bacteria, *Erwinia carotovora* (Er), Medkleb (Med) and *Rhizobium leguminosarum* (Rhiz) was measured via the quantity of reduced sugar in solution, following incubation of culture filtrate with polygalacturonic acid. Units of polygalacturonase were quantified in terms of colour change via extrapolation from the standard curve. Med and Er filtrate contained 30 and 46 units of polygalacturonase per ml respectively. Rhiz filtrate did not contain any polygalacturonase. C) Units of polygalacturonase contained in bacterial filtrate. Units of polygalacturonase per ml are represented on the y-axis. 1000ul aliquots of bacterial filtrate processed from three different species of bacteria are represented of the x-axis. Bacteria were assessed for the presence of polygalacturonases using a standard curve of galacturonic acid. The upper and lower hinges of boxplots represent the first and third quartiles of enzyme concentrations in filtrate, calculated from three technical replicates. *Erwinia carotovora* was the highest producer, with an average of 45.7 units/ml and Medkleb produced polygalacturonase with an average of 29.7 units/ml. *Rhizobium leguminosarum* did not produce any polygalacturonase.

#### (iii) Predicted metabolic functions of Medkleb

KEGG (Kanehisa et al 2016) was used to produce a list of gene functions for chromosomes and plasmids associated with all *K. oxytoca* bacteria in the Medkleb group (Table S5). The dataset was filtered to produce three smaller lists: 1) Atypical -gene functions unique to Medklebn, 2) Absent -gene functions present in all genomes analysed other than Medkleb, 3) Duplicated – gene functions encoded in multiple copies by Medkleb but not by any other genome analysed The Medkleb genome contained 21 atypical functional orthologues (Figure S7), and of these, 16 were chromosomally derived and five located on plasmids. The atypical genes included two putative transposases, enzymes involved in various modes of metabolism (e.g., *mtlA, ACADSB, egsA, FAAH2*) and four transport protein genes (e.g. *natA & gatC*). When compared to conspecifics, Medkleb has 35 absent gene functions, seven of which may indicate adaptation to the relatively innoxious medfly gut (Figure S8). The largest cluster of absent genes was associated with copper resistance (*copB, cusS, cusR, cusC, cusF)* and genes that degrade arsenic (*arsB*) and nitriles (*nthA*) were also missing. The Medkleb genome codes for multiple copies of 11 functional orthologues, that other *K. oxytoca* bacteria in the Medkleb group retain in single copy at most (Figure S9). Medkleb’s duplicated genes may have the potential to promote mutualistic phenotypes for the medfly, as they syntheise amino acids (*proC, trpA*) and metabolise beneficial nutrients such as fatty acids (*ACAT*) and citrate (*citE*).

#### (iv) Secondary metabolites

The Medkleb genome and plasmid sequences were interrogated using antiSMASH 6.0 beta (Blin et al 2019) for the presence of novel secondary metabolites. Chromosomes of all *K. oxytoca* members of the Medkleb group (other than KONIH2) coded for three common secondary metabolite clusters: 1) non-ribosomal polypeptide synthetase (30% similarity to turnerbactin), 2) thiopeptide antibiotic (14% similarity to O-antigen), 3) Ribosomally synthesized and post-translationally modified peptides (RiPPs). The Medkleb chromosome contained two unique secondary metabolite clusters that were not present in any other analysed *K. oxytoca* chromosomes (i) a butyrolactone, a signalling molecule utilised by *Streptomyces* bacteria to to regulate antibiotic production and cell cycle processes (Takano 2006; Kitani et al 2011), and (ii) an N-acyl amino acid cluster, common to soil dwelling bacteria and involved in cell-to-cell communication (Craig et al 2011; Battista et al 2019) (Table 2). Plasmid mkp4 was also predicted to code for cloacin and colicin bacteriocins that were not coded by any other *K. oxytoca* strain in the Medkleb group. The cloacin cluster encoded on mkp4 contains two mobile elements (MKleb_5887, MKleb_5890) and is generally considered to be toxic for *Klebsiella* bacteria (de Graaf et al 1969).

**Table 2.** Unique features of the Medkleb genome in comparison to conspecifics.

| Unique feature | Identified via | Possible mutualistic function |
| --- | --- | --- |
| *LCB18* | progressiveMauve analysis of Medkleb and three conspecifics | Codes for genes related to molybdopterin biosynthesis and several proteins with unknown function. These genes may confer fitness to the medfly. |
| *GI1* | Inversion of GC skew | Codes for genes related to molybdopterin biosynthesis and several proteins with unknown function. These genes may confer fitness to the medfly. |
| *Atypical metabolic genes* | Comparison of KEGG gene functions between Medkleb group members. | *mtlA*, *ACADSB*, *egsA* and *FAAH2* expand the range of nutrients available to the medfly. |
| *Absent sensory genes* | Comparison of KEGG gene functions between Medkleb group members. | The medfly gut may be relatively non-toxic allowing Medkleb to survive without genes that detect and metabolise certain chemical threats in its environment (*copB*, *cusS*, *cusR*, *cusC*, *cusF*, *arsB*, *nthA*). |
| *Duplicated genes* | Comparison of KEGG gene functions between Medkleb group members | Duplication of 11 genes associated with biosynthesis of amino acids and breakdown of essential nutrients. These genes may confer fitness to the medfly by providing access to extra nutritional resources |
| *Butyrolactone biosynthesis* | antiSMASH 6.0 beta | Possible regulation of antibiotic products (Takano 2006; Kitani et al 2011) |
| *N-acyl amino acid biosynthesis* | antiSMASH 6.0 beta | An important family of endogenous signalling molecules in which an amide bond covalently links an amino acid to the acyl moiety of a long-chain fatty acid. Primarily involved in cell-to-cell communication (Craig et al 2011; Battista et al 2019_ |

### 5. Searching for symbiotic signatures in the Medkleb genome

#### (i) Genome size and GC content

At 5_825_435bp, the Medkleb genome was the smallest in the ‘Medkleb group’ and 1.6 standard deviations smaller than the mean genome size of the clade (n=13) 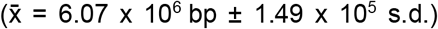. However, we conclude this is not diagnostic of symbiotic transition as the genome of free-living *K. michiganensis* strain M1, was only 0.7% larger than Medkleb (5.86 × 10^6^ bp). Medkleb’s GC content (56.03%) was also high, even for bacteria with a free-living life history (Moran et al 2008) (Table 1). We conclude that there were no obvious diagnostic signatures of a strong host-association.

#### (ii) Local genomic rearrangement

The Medkleb genome was aligned, using progressive Mauve (Darling et al 2010) to three closely related strains of *K. oxytoca* (Figure S5). This analysis revealed the presence of 21 local colinear blocks (LCBs) of conserved DNA in all four genomes (Figure S5). These LCBs were not uniformly distributed between the genomes. *K. oxytoca* strains AR0147 and CAV1374 both contained inversions between LCBs 11-16 and strain KONIH2 contained several instances of translocation and inversion. However, despite inversion and translocation events, nucleotide sequence in all LCBs other than LCB18 was very highly conserved. The annotation of Medkleb LCB18 (≈nts 4.97 × 10^6^ ∼ 5 × 10^6^) predicts 25 coding sequences in total, including 13 “hypothetical proteins”, 2 DNA helicases (Mkleb_4576/4577), 2 methyltransferases (Mkleb_4579/4597) and an anti-restriction protein (Mkleb_4581). LCB18 may have been recently acquired as it also contains genes associated with horizontal transfer including an integrase (Mkleb_4574), a mobile element protein (Mkleb_4594) and a prophage protein (Mkleb_4585). In addition, two hypothetical proteins encoded by LCB18 (MKleb_4591 & Mkleb_4592) may be virulence factors, as they are predicted by TMHMM 2.0 {Krogh, 2001) to encode N-terminal peptides. However, there is as yet no definitive evidence that LCB18 contains genes that might confer fitness to the medfly (Table 2).

The Medkleb genome carries an inversion of GC skew on the leading strand (>50% C; region GI1) indicating a putative horizontal gene transfer event. GI1 is thought to have been transferred from a plasmid, as it codes for *TraY* which facilitates plasmid conjugal transfer (Nelson et al 1995). Though plasmid integration into host chromosome is common (Dobrindt et al 2004; Bire and Rouleux-Bonnin 2012), GI1 does not appear to have integrated from any of the Medkleb plasmids (mkps 1-4) (Table 2). In total GI1 is predicted to code for 25 proteins including Bll0873, first sequenced in *Bradyrhizobium diazoefficiens*, a bacterium that is a known nitrogen fixing symbiont of legumes (Kaneko et al 2002). Interestingly, both GI1 (Mkleb_0586) and LCB18 (Mkleb_4596) code for genes predicted to facilitate molybdopterin biosynthesis, which could have the potential to benefit the medfly host, as this group of co-factors aid nitrate reduction (Moreno-Vivián et al 1999).

According to KEGG analysis, the Medkleb genome contains coding sequences for 21 atypical functional orthologue genes that are not encoded in the genomes of any other closely related *Klebsiella* bacteria in the Medkleb group (Table 2). Some of these are potentially mutualistic functions such as sugar metabolism (*mtlA*) and nitrogen fixation (*nifZ*). Medkleb also has duplicates of 11 functional orthologues that *K. oxytoca* generally retains in only single copy. These duplicated functional orthologues do not cluster by genomic location. Several of these duplicated gene functions have strong mutualistic potential such as the biosynthesis of amino acids (*proC, trpA*) and breakdown of essential nutrients (*cite, pydC*). In contrast, Medkleb is also missing some clusters of genes that are present in closely related bacteria, e.g. for specific enzymes related to copper resistance (*copB, cusS, cusR, cusC, cusF*) and phosphonate transport (*phnC, phnD, phnE)*.

## Discussion

### The medfly microbiome

In this study, we report the use of laboratory and field-reared adult medflies to characterise key features of the bacterial microbiome of this important agricultural pest. Using analyses of 16S rRNA gene amplicon sequencing of culturable and non--culturable members of the gut microbiome, we found that overall species richness was fairly stable between the laboratory-reared flies raised on different diets and the wild medfly samples. Direct comparisons of beta diversity indicated that larval diet, rather than exposure to antibiotics or wild vs lab rearing was the primary driver of microbial diversity of gut microbiomes. Wild flies obtained from distinct geographical regions and different hosts and flies reared on different substrates and exposed to antibiotic cocktails in the laboratory, contained largely the same bacterial families. The data suggest that although medflies are highly polyphagous, they have a stable microbiome that is dominated by the bacterial family *Enterobacteriaceae*, including a putative symbiont *Klebsiella* spp. These findings are consistent with previous analyses of medfly microbiomes made using culture-based methods (Behar et al 2005; Behar et al 2008a; Behar et al 2008b; Behar et al 2009a). The picture may be more complex, however, as one recent next-generation sequencing study did not isolate *Klebsiella* in medfly microbiomes from wild populations (Malacrinò et al 2018), and another identified possible geographic or host-genetic structure links with bacterial dominance (with samples from Greece being dominated by *Klebsiella* spp) (Nikolouli et al 2020).

### Genome sequencing and analysis of the putative Medkleb gut symbiont

We obtained a fully sequenced Medkleb genome and both 16S rRNA gene and ANI phylogenies identified this as a *Klebsiella* species (Stackebrandt 2006; Kim et al 2014). Medkleb was *pehX* positive when analysed with PCR, and a producer of pectinolytic enzymes, both of which support its identification as *K. oxytoca* (Kovtunovych et al 2003; Saha et al 2013).

Genome size and GC content are generally reduced when a bacterium adopts a facultative lifestyle (McCutcheon and Moran 2011). We found that, although the Medkleb genome was the smallest of all *Klebsiella* bacteria in the ‘Medkleb group’, the free-living *K. michiganensis* strain M1 was only 40 kb (0.7%) larger. The Medkleb GC content was also the second highest in the ‘Medkleb group’, again counter to what is expected for a facultative mutualist. A strongly symbiotic transition is generally associated with an increased mutation rate, which causes facultative symbionts to occupy extended branches of phylogenetic trees (McCutcheon and Moran 2011). Though Medkleb was slightly distinct from the three bacterial species most closely related to it, it was not markedly so.

We also used high resolution comparative genetic techniques to test for signals of putative mutualism. These analyses were designed to test: (i) If Medkleb possessed genetic loci/functions that were absent from closely related *Klebsiella* bacteria and possessed an obvious mutualistic capacity for the fly; (ii) If Medkleb was lacking genetic loci/functions found in all closely related free-living *Klebsiella* spp., that would be considered necessary for life in varied environments. Set against this is that gene loss may be unpredictable in the early stages of facultative mutualistic transitions (Moran and Bennett 2014).

Comparative genomics highlighted that the Medkleb genome contained two discrete regions (GI1 and LCB18) which may have been acquired comparatively recently and are not typically found in *K. oxytoca*. Both GI1 and LCB18 contain genes associated with horizontal transfer and encode several unannotated hypothetical proteins with mutualistic potential. Medkleb was demonstrated to be functionally pectinolytic and global gene function analyses highlighted several more atypical functions were encoded exclusively by Medkleb in comparison to conspecifics. These atypical genes are dispersed throughout the genome which suggests that they are ancestral and not recently inherited. Some of these genes are predicted to encode metabolic enzymes that could allow the medfly to utilise otherwise unattainable nutrients. In addition, Medkleb has several duplicated genes that are randomly distributed throughout the genome which could provide the medfly with beneficial nutrients and amino acids. Together, these genomic signatures suggest that Medkleb is a symbiont that improves host fitness through increased access to vital nutrients. If substantiated, this could help to partially explain the success of the medfly as a generalist pest. However, potentially mutualistic traits are not evidence of mutualism itself and many signals that are commonly associated with facultative mutualism are not evident in the Medkleb genome. Therefore there is insufficient evidence, so far, to definitively classify Medkleb as a facultative mutualist of the medfly.

*Klebsiella* bacteria and the medfly appear to co-occur almost universally (Behar et al 2009b; Leftwich 2012). Therefore, Medkleb might provide a platform for paratransgenic control of the medfly. This study is one of only a handful of comprehensive characterisations of the bacterial symbionts of the tephritid fruit fly *C. capitata* using culture-independent methods, and the first to characterise the genome of a putative symbiont. Future studies should sample all developmental stages of the medfly, and sample more intensively to establish the potential for phylosymbiosis. Complementing this should be experimental studies to establish localisation of *Klebsiella* within the medfly gut, to determine transmission mechanisms and assess metabolic communication and host benefit (Leftwich et al 2020).

## Data Deposition

Sequencing data has been submitted to the NCBI Sequence Read Archive and is available under BioProject ID PRJNA638617.

## Acknowledgements

This work was funded by the Biotechnology and Biological Sciences Research Council Research (Grant BB/K000489/1 to T.C., P.T.L., and M.I.H. and PhD studentship to M.D). We also thank Kostas Voudouris, Romisa Asadi & Martha Koukidou for their help in collecting samples for analysis.

