## Supplemental Files for "Characterisation of the symbionts in the Mediterranean fruitfly gut"

### Supporting Information

### 1. Fly rearing for tests of the effects of different larval diets on the adult microbiome

Three independent biological replicates of each diet treatment were maintained under strict allopatric conditions. All experiments and culturing were conducted at 25°C, 50% relative humidity, on a 12:12 light dark photoperiod. Adults emerging from each replicate were maintained in groups of roughly 30 males and 30 females in plastic cages (11 cm × 11cm × 10 cm). Adults from all lines received the same standard adult diet (*ad libitum* access to sucrose-yeast food; 3:1 w/w yeast hydrolysate/sugar in water). Approximately 500 eggs were placed on 100 mL of each of the appropriate larval diets (Table S1) in a glass bottle. When third instar larvae started to "jump" from the larval medium, the bottles were laid horizontally on sand and pupae allowed to emerge for seven days. Pupae were then sieved from the sand and held in 9 mm petri dishes until adult eclosion. Adult samples were collected at 5 and 10 days post eclosion. The effect of each larval diet was tested in the presence and absence of antibiotics. Antibiotic treatments comprised 100µg/ml kanamycin, 200 µg/ml ampicillin, 200 µg/ml streptomycin, 50 µg/ml chloramphenicol, 100µg/ml apramycin, 100 µg/ml hygromycin and 200 µg/ml tetracycline.

### 2. Taxonomic analysis of 16SrDNA sequences of medkleb

The EZBioCloud (Yoon et al., 2017) was searched for sequences homologous to Medkleb's putative 16S_rDNA sequence from nucleotides 341370-342821 (mk16S). The 16S sequence with greatest homology to mk16S belonged to the W14 strain, which had been classified as *K. michiganensis*. Medkleb's predicted species classification was *K. oxytoca,* hence near identical 16S homology with *K. michiganensis* was unexpected, although it is recognised that 16S sequence similarity alone does not confirm species identification (Tindall et al., 2010). As recommended by Tindall et al. (2010), homology between mk16S and 16S sequences that had been classified as either *K. oxytoca* or *K. michiganensis* was calculated using EZBioCloud (Yoon et al., 2017) but this also failed to distinguish Medkleb as either species. mk16S was compared with 40 16S sequences classified as either *K. oxytoca* or *K. michiganensis* and was found to be between 98.5 and 99.93% related to all strains analysed. Stackebrandt (2006) suggest that >98.7% similarity should be the threshold at which 16S sequences are considered to be conspecific but some authors have suggested thresholds need to be as high as 99.5% (Janda et al., 2007). In conclusion, Medkleb could not be specifically characterised as either *K. oxytoca* or *K. michiganensis* via pairwise analysis of its 16S sequence, as according to current taxonomic standards (Janda et al., 2007; Stackebrandt, 2006), the *K. oxytoca* and *K. michiganensis* strains analysed here were themselves conspecific.

**Commented [MD1]:** Medkleb's putative 16S sequence (nucleotides 341370-342821;mk16S)

To classify Medkleb's species with improved resolution, a comprehensive 16S phylogenetic analysis (Figure S3) was carried out. RefSeq 16S sequences used in the analysis had been classified as either *K. oxytoca, K. michiganensis* or *K. pneumoniae* (Quast et al., 2013), and a single strain of *Pseudomonas aeruginosa* was used as an ancestral root for the phylogeny. *K. pneumoniae* sequences were included as a control, as this species is closely related to *K. oxytoca* (Kovtunovych et al., 2003) but distant enough to be distinguished phylogenetically. Medkleb, *K. oxytoca* and *K. michiganensis* sequences formed one homogenous group and *K. pneumoniae* formed an outgroup, suggesting that Medkleb is both *K. oxytoca* and *K. michiganensis* and hence that the species dichotomy is artificial, as suggested above. However, even though 16S phylogenetic and pairwise analyses failed to delineate *K. oxytoca* from *K. michiganensis*, these species should be considered distinct, as DNA-DNA hybridisation between them is <70% (Saha et al., 2013).

**Supplementary figures**

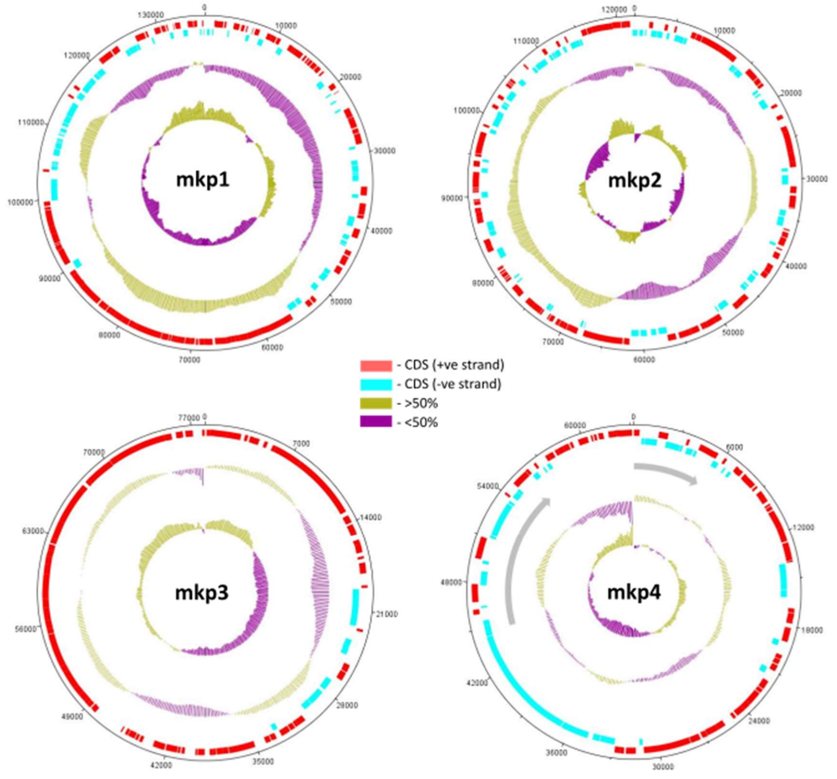


**Figure S1. Circular maps of plasmids - mkps 1-4.** For all maps: 1) red sectors on track one represent gene coding sequences on the positive strand, 2) light blue sectors on track two represent gene coding sequences on the negative strand, 3) track three represents GC content; regions with >50% GC content are mustard and regions with <50% GC content are purple, 4) track four represents GC skew; regions with a >50% G's are mustard and regions with <50% G's are purple. Interesting individual plasmid features include: 1) mkp4 contains two secondary metabolite clusters indicated by grey arrows, 2) mkps 1, 3 and 4 exhibit clear coding bias.

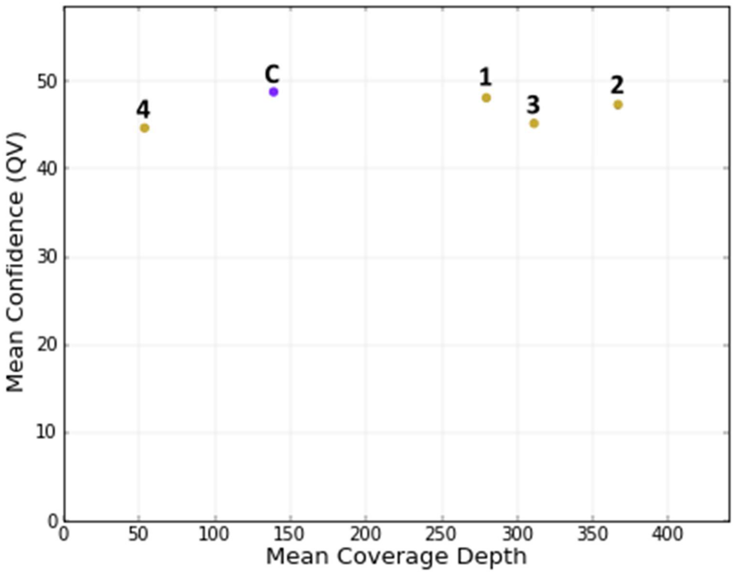


**Figure S2. Contig coverage vs quality value for Medkleb sequencing contigs.** The mean coverage depth for the Medkleb chromosome (purple dot) was 138.7 reads per base, and mean QV was estimated at 48.9. The four plasmids (yellow dots) had mean coverage depths and QV's of: mkp1) 279.6 and 48; mkp2) 366.4 and 47.3; mkp3) 310.8 and 45.3; mkp4) 53.6 and 44.8. Coverage was relatively high in the cases of mkp1-3, which was as predicted, as plasmids are often retained in high copy number.

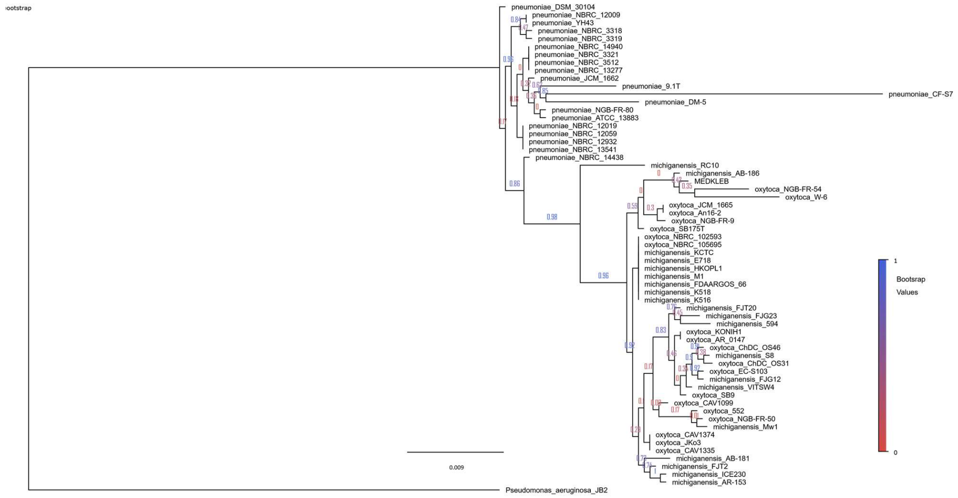


**Figure S3. Phylogeny showing the evolutionary relationship of 16S genes belonging to Medkleb and three species of the *Klebsiella* genus.** The phylogeny was created with the Silva ACT service (Pruesse et al., 2012), the FastTree2 maximum likelihood program (Price et al., 2010) and rooted with *Pseudomonas aeruginosa* strain JB2. The scale bar represents substitutions per site. Bootstrap values are represented at all nodes and are colour coded according to confidence level (blue = 1 and red = 0). 16S sequences of strains that had been identified as *K. pneumoniae* demonstrated clear evolutionary divergence and segregated definitively from *K. oxytoca* and *K. michiganensis* strains. 16S sequences identified as *K. oxytoca* and *K. michiganensis* formed one homogenous group, with some species level clustering but no reliable pattern of distribution. Species identification of Medkleb was not possible based on this phylogeny, as it segregated within the *K. oxytoca, K. michiganensis* group.

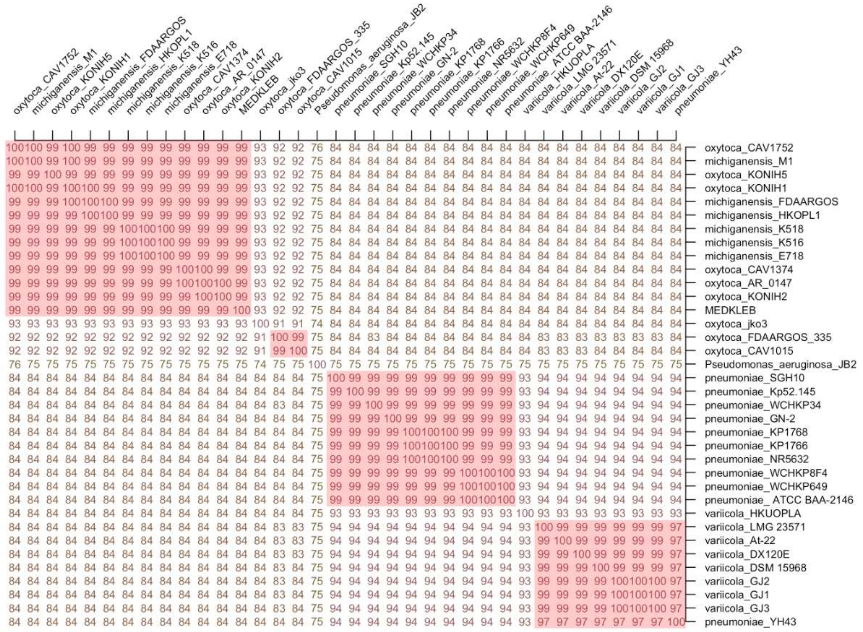


**Figure S4. Average nucleotide identity matrix for 35 strains of *Klebsiella* bacteria.** The matrix was created using the ANI calculator (Figueras et al., 2014), with *Pseudomonas aeruginosa* strain JB2 selected as the outgroup. Not including the *Pseudomonas* outgroup, six distinct *Klebsiella* species were identified by the ANI analysis. Isolates with >95% ANI identity are considered conspecific and are highlighted pink. The “Medkleb group” is located at the top left of the matrix.

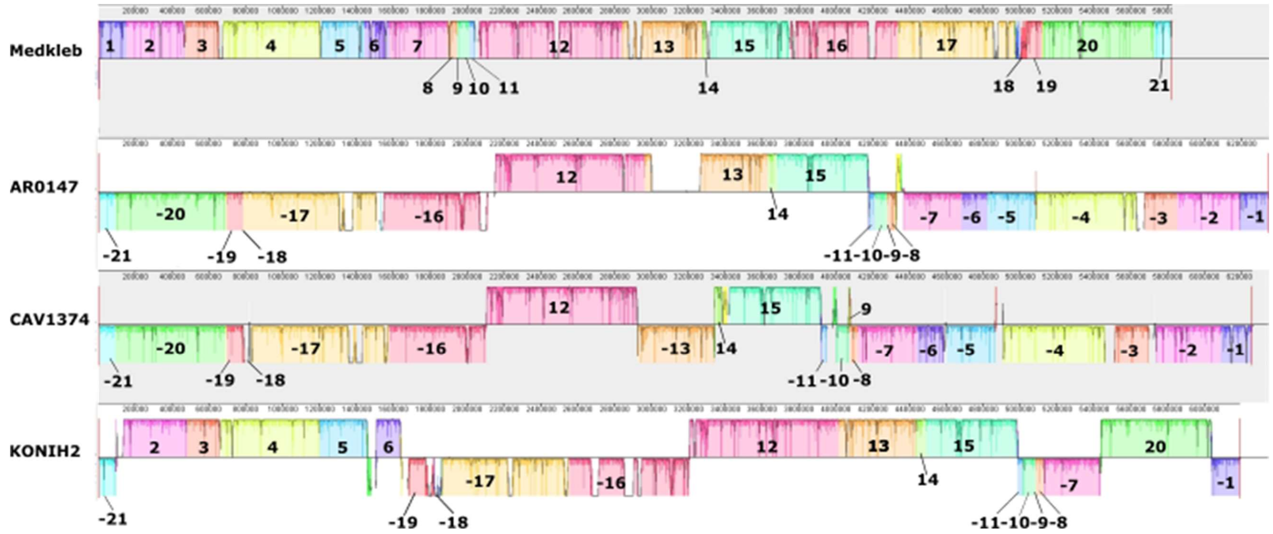


**Figure S5 Synteny plot of Medkleb and three closely related bacteria.** Medkleb was positioned phylogenetically in a clade with three *K. oxytoca* bacteria (strains AR0147, CAV1374 and KONIH2) with ANI analysis (Konstantinidis et al., 2005). progressiveMauve (Darling et al., 2010) was then used to assess synteny between Medkleb, AR0147, CAV1374 and KONIH2. This analysis, which is represented graphically above, uncovered 21 conserved local colinear blocks (LCBs) (Darling et al., 2010) of nucleotide sequence that were shared between all genomes analysed. Block height corresponds the degree of regional sequence conservation. When compared to Medkleb, sequence conservation in block 18 was low for strains AR0147, CAV1374 and KONIH2. There were several instances of LCB inversion between strains but inverted LCBs were not low in height, which indicated that nucleotide sequence was conserved.

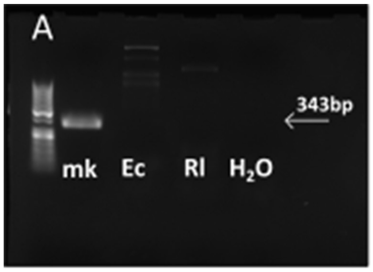


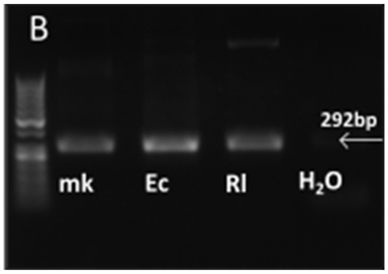


**Figure S6. PCR amplifications of *pehX* and 16S rRNA genes.** Both gels were run with three samples of DNA and a water control. A) PCR amplification of the *pehX* gene produced the expected 343bp product from Medkleb (mk). Neither *Erwinia carotovora* (Ec) nor *Rhizobium leguminosarum* (Rl) amplified this product, but both (Ec in particular) appeared to amplify products larger than 343bp. B) Control reactions: all three species of bacteria produced the expected 292bp 541F-806R amplicon following PCR amplification of 16S, which verified the quality of DNA and reagents used in the PCR assays.

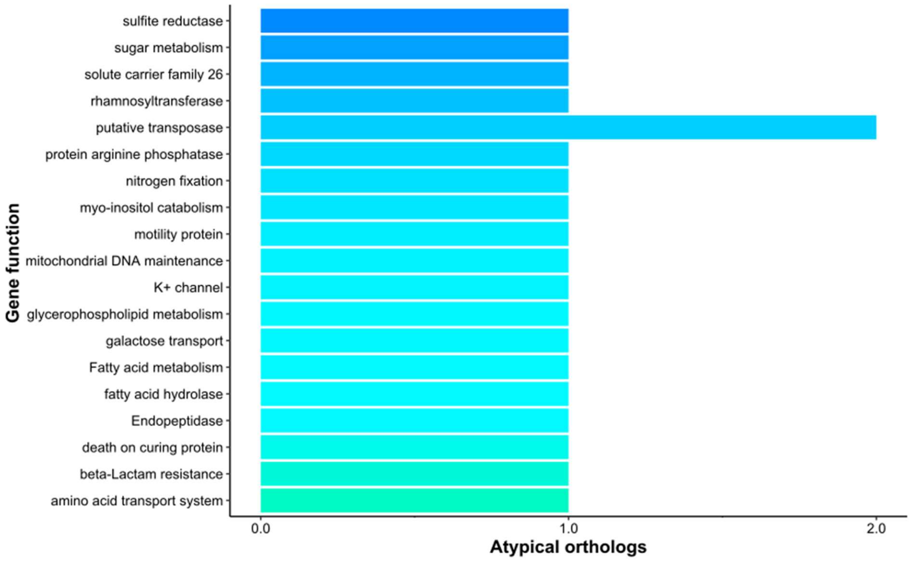


**Figure S7. The functions of genes present in the Medkleb genome but absent from all other *K. oxytoca* bacteria in the Medkleb group.** The functions of atypical genes that were present in the Medkleb genome but absent from all remaining *K. oxytoca* genomes analysed are represented on the y-axis. Each atypical function is denoted by a discrete colour. The number of atypical genes providing each function is represented on the x-axis. Medkleb codes for 21 atypical genes with 20 different functions. Broadly, these genes were predicted to be involved in metabolism, transport and gene transposition.

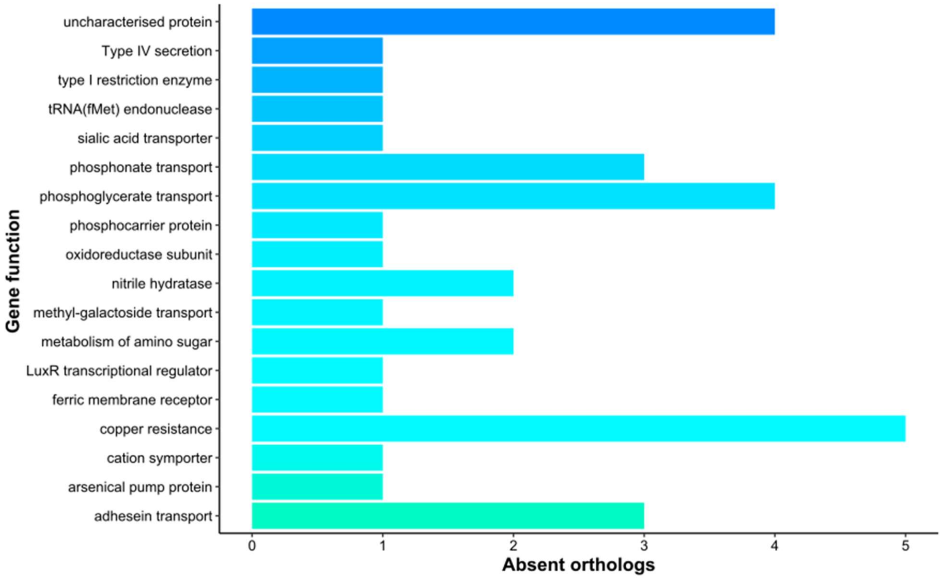


**Figure S8. The functions of genes that are absent from the Medkleb genome but present in all other *K. oxytoca* bacteria in the Medkleb group**. The functions of genes that were absent from the Medkleb genome but present in all other *K. oxytoca* genomes analysed are represented on the y-axis. Each absent function is denoted by a discrete colour. The number of absent genes associated with each function is represented on the x-axis. Medkleb did not code for 35 genes that were present in every other genome analysed and these were associated with 17 discrete functions. Loss of function occurred in clusters of genes relating to the detoxification of xenobiotic compounds.

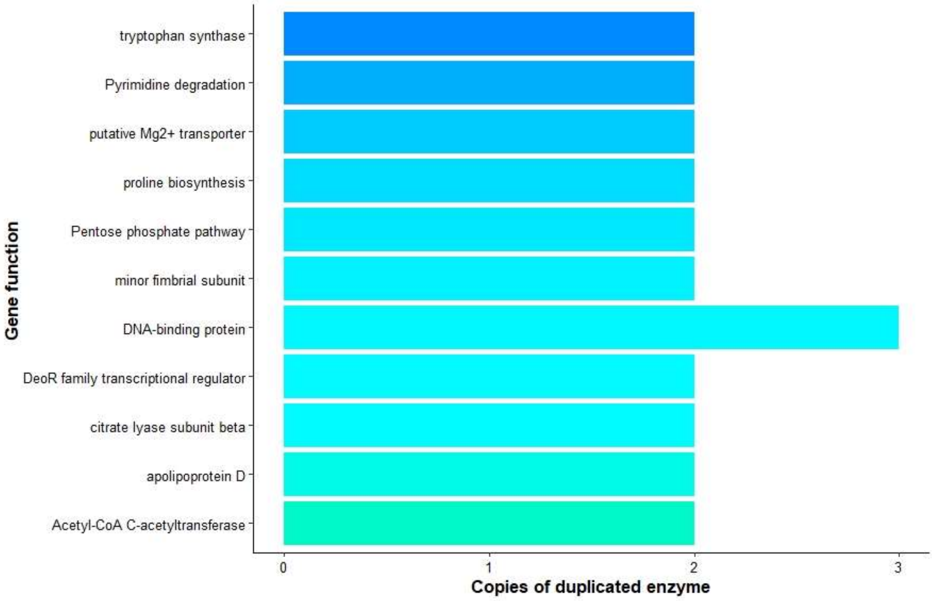


**Figure S9. Gene functions duplicated by Medkleb but not duplicated by any other *K. oxytoca* bacteria in the Medkleb group**. Genes functions held in multiple copies by Medkleb but not by any other *K. oxytoca* genome analysed are represented on the y-axis. Each duplicated function is denoted by a discrete colour. The number of gene copies associated with each function is represented on the x-axis. Medkleb has duplicated 11 genes that are either absent or held in single copy by all other genomes analysed. Duplicated functions include the biosynthesis of amino acids and degradation of citrate.

**Supplementary tables**

**Table S1. Larval diets used for laboratory rearing**

| | | | |
|---|---|---|---|
| **Sucrose** | Water | 1000 | ml |
| | Agar | 15 | g |
| | Sucrose | 30 | g |
| | Yeast | 50 | g |
| | Propionic Acid | 5 | ml |
| **Low Protein (Sucrose)** | Water | 1000 | ml |
| | Agar | 15 | g |
| | Sucrose | 30 | g |
| | Yeast | 30 | g |
| | Propionic Acid | 5 | ml |
| **Starch** | Water | 1000 | ml |
| | Agar | 15 | g |
| | Starch | 30 | g |
| | Yeast | 50 | g |
| | Propionic Acid | 5 | ml |
| **Glucose** | Water | 1000 | ml |
| | Agar | 15 | g |
| | Glucose | 30 | g |
| | Yeast | 50 | g |
| | Propionic Acid | 5 | g |

**Table S2. Contig derivation and GC content.** mlplasmids (Arredondo-Alonso et al., 2018) predicted that the largest of the five Medkleb contigs was chromosomal, with a posterior probability (p(θ\X)) of 0.991. mkp2 and mkp5 were predicted to be plasmid-derived, with posterior probabilities of 0.962 and 0.977 respectively. The posterior probabilities of mkp3 and mkp4 being plasmid-derived were less powerful but still robust, at 0.864 and 0.882 respectively. mkp2 and mkp4 both had GC content comparable to that of the putative chromosomal contig, suggesting they had arisen as a result of a recent horizontal transfer.

| **contig** | **length** | **p(θ\X)** | **Prediction** | **GC**(%) |
|---|---|---|---|---|
| 1 | 5825435 | 0.009 | Chromosome | 56.03 |
| 2 | 136402 | 0.962 | Plasmid | 55.18 |
| 3 | 122124 | 0.864 | Plasmid | 50.4 |
| 4 | 78046 | 0.882 | Plasmid | 56.68 |
| 5 | 63257 | 0.977 | Plasmid | 50.65 |

**Table S3. Synopsis of the Medkleb genome.** The Medkleb genome was 5867451 nt in length and coded for a total of 5388 genes, at a coding density of 0.941 genes per kb.

| | |
|---|---|
| **Length (nt)** | 5825435 |
| **GC content (%)** | 56.03 |
| **CDS (+ve strand)** | 2541 |
| **CDS (-ve strand)** | 2847 |
| **Overall coding sequence (%)** | 87.8 |
| **Gene density (per kb)** | 0.941 |

**Table S4. Comparison of plasmid genomes.** mkps 1-4 varied in length by over 100%, and GC content varied by 6.3%. Plasmid 4 was relatively small but had the greatest gene density, utilising approximately 4% more nucleotide sequence for protein coding than other plasmids.

| | **mkp1** | **mkp2** | **mkp3** | **mkp4** |
|---|---|---|---|---|
| **Length (nt)** | 136402 | 122124 | 78046 | 63257 |
| **GC content (%)** | 55.2 | 50.4 | 56.7 | 50.7 |
| **CDS (+ve strand)** | 105 | 95 | 75 | 62 |
| **CDS (-ve strand)** | 70 | 64 | 8 | 42 |
| **Overall coding sequence** | 80.3 | 81.1 | 81.2 | 85.3 |
| **Gene density (per kb)** | 1.28 | 1.31 | 1.06 | 1.64 |
| **Annotated plasmid CDS** | 12 | 3 | 18 | 12 |
| **Mobile element protein** | 21 | 21 | 1 | 5 |

**Table S5 Comparison of gene functionality for *K. oxytoca* bacteria in the Medkleb group.** The total number of gene functions was fairly consistent in this group, at approximately 2900 per strain. In terms of atypical functions, Medkleb had a relatively large complement, but fewer than CAV1752. The Medkleb genome had more absent gene functions than was found for conspecifics.

| | **Total gene functions** | **Atypical functions** | **Absent functions** |
|---|---|---|---|
| **Medkleb** | 2857 | 21 | 35 |
| **AR0147** | 2904 | 12 | 4 |
| **CAV1752** | 2882 | 24 | 5 |
| **CAV1374** | 2944 | 11 | 6 |
| **KONIH1** | 2917 | 18 | 3 |
| **KONIH2** | 2912 | 13 | 10 |
| **KONIH5** | 2882 | 11 | 5 |
